# A Hybrid Model for the Effects of Treatment and Demography on Malaria Superinfection

**DOI:** 10.1101/476523

**Authors:** John M. Henry

## Abstract

As standard mathematical models for the transmission of vector-borne pathogens with weak or no apparent sterilizing immunity, Susceptible-Infected-Susceptible (SIS) systems such as the Ross-Macdonald equations are a useful starting point for modeling the impacts of interventions on prevalence for diseases that cannot superinfect their hosts. In particular, they are parameterizable from quantities we can estimate such as the force of infection (FOI), the rate of natural recovery from a single infection, the treatment rate, and the rate of demographic turnover. However, malaria parasites can superinfect their host which has the effect of increasing the duration of infection before total recovery. Queueing theory has been applied to capture this behavior, but a problem with current queueing models is the exclusion of factors such as demographic turnover and treatment. These factors in particular can affect the entire shape of the distribution of the multiplicity of infection (MOI) generated by the superinfection process, its transient dynamics, and the population mean recovery rate. Here we show the distribution of MOI can be described by an alternative hyper-Poisson distribution. We then couple our resulting equations to a simple vector transmission model, extending previous Ross-Macdonald theory.

## 1 Introduction

Malaria continues to have a significant impact on the world, and is responsible for over 400,000 deaths in 2015 alone [4]. Over the past century a repertoire of interventions has been developed to prevent its spread and reduce its impact on those afflicted, but the parasite continues to adapt and persist. Because malaria dynamics and control are so complex, mathematical and statistical models play a significant role in the strategic implementation of these existing tools to maximize their impact. One of the historically difficult aspects of malaria and other macroparasitic diseases to model is the process of superinfection, wherein multiple distinct cohorts of pathogens infect the host simultaneously. The standard SIS system usually used to model similar disease transmission using a constant recovery rate from a single infected class does not capture this type of heterogeneity, and may therefore be a poor approximation to the real system.

There has been a long history of modeling a number of disease states to track the proportion of individuals with *n* infections, each of which is still susceptible to further infection [2, 3, 8]. Care must be taken in modeling the distribution of the multiplicity of infection (MOI), that is the fraction of the population that has exactly *n* “broods” or cohorts of parasites. This distribution strongly affects the population mean recovery rate, which can be seen as a weighted sum of the rate at which individuals are recovering from *n* infections weighted by the proportion of individuals with *n* infections. The shape of this distribution can be profoundly altered by factors such as seasonality and case management, which in turn can have considerable effects on the mean rate of recovery for the population and the prevalence of the disease.

Under George Macdonald’s original assumption that infection status is no barrier to reinfection, MOI can be modeled as an M/M/*∞*queue [2, 8]. Though Macdonald’s written description of the system was correct, he mistakenly presented it mathematically as an M/M/1 queue [3]. This system vastly overestimates an individual’s time spent infected, with the average becoming infinite if the force of infection exceeds the rate of recovery. Macdonald’s model and its description generated some confusion, substantial discussion, and a series of approximating models [10, 12].

However all of this analysis has historically been focused on the original M/M/*∞*model, whose assumptions include a strong steady state condition on the population wherein individuals experience no demographic turnover or external forces that influence their susceptibility or infection status. Critically, finite lifespans and treatment will affect the distribution of MOI. Including their interactions ad hoc in ordinary differential equation (ODE) models may lead to inconsistent transition rates, and even inconsistent equilibria, when compared to the original assumptions of the stochastic model.

For models in which infections are assumed to clear independently, the mean population recovery rate is dependent on the proportion of individuals with a single infection - therefore the distribution of MOI strongly influences the recovery rate. Excluding demographics and allowing for variable force of infection (FOI, here denoted *λ*) allows hybridization of the transmission system given the distribution is Poisson, and it was modeled elegantly by Nåsell with two relatively simple ODEs [9]. However, introducing demography and treatment requires a different approach as they can fundamentally alter the process generating the distribution of MOI.

Here we consider the system with demography and treatment, with and without transient chemo-protection, and present a new way of modeling the statics and dynamics of the distribution of MOI and its nonlinear effects on the population mean recovery rate and resulting prevalence. In addition, we show that this distribution is stable in the sense that any initial distribution with finite mean converges in distribution to the stationary distribution. We then show the impact of an extended protective period post-treatment, which has the effect of inducing a zero-inflation in the MOI distribution. Using these results and their resulting dynamics, we then extend the Ross-Macdonald model to include this heterogeneity and explore the effects of mortality and treatment on prevalence under these assumptions.

## 2 Model with Demography and Treatment, without Chemoprotection

### 2.1 Model Description and Setup

Consider a Poisson process *X*_t_(*λ*) of new infections in an individual person with intensity *λ*, the force of infection (FOI). We will assume *λ* may be time-varying, but it is independent of the current number of infections as per Macdonald’s original assumption. In addition, we will have a second Poisson process *Y*_t_(*rX*_t_) of recovery with an intensity *rX*_t_. Again, we make no assumption about r aside from it being independent of *X*_t_ or *Y*_t_. Then the M/M/*∞* queue can be represented by a third Poisson process, defined to be the difference of the two: *Z*_t_ := *X*_t_ − *Y*_t_. This process counts the current number of infections in an individual, and its range is on the non-negative integers. To add demography and treatment, which both act to decrease from the infected classes and increase the susceptible class, we need additional Poisson processes *D*_t_(*δ*) and *T*_t_(*τ)*, where *δ* and *τ* are respectively the mean birth/death rate (assumed equal to maintain constant population sizes) and the mean treatment rate. Treatment only acts on those with at least one infection, and because births are assumed to replace deaths, demography also has a net effect of removing infected individuals and introducing susceptible individuals. Therefore for brevity we can replace the two processes with a single Poisson process *M*_t_ with rate *µ*, where *µ* := *δ* + *τ* is the expected rate at which either death or treatment occurs.

The master equations for the system can be built by considering the state space in Figure 1:

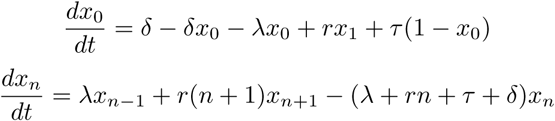

which we can rewrite using the combined treatment and demographic process with parameter *µ*:

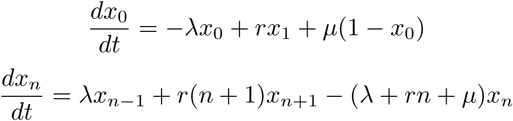

**Figure 1:**
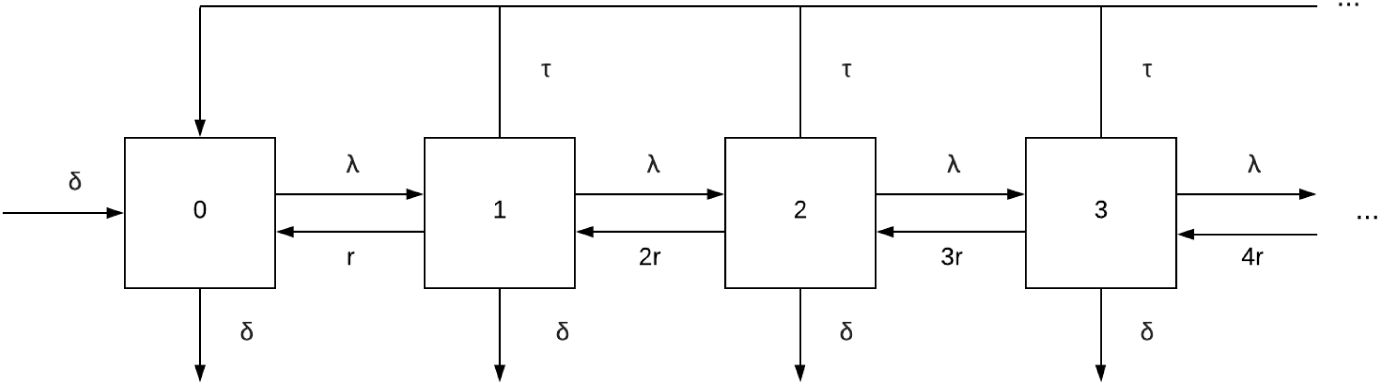
State space of human infections without chemoprotection. Numbers indicate the number of infections in an individual and the letters denote transition rates between states.

As this is an upper triangular infinite system of linear ODEs, it can’t be solved iteratively; instead we will introduce a generating function of the form

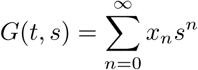

which encodes the infinite variables as coefficients in its power series, simplifying further analysis. We can directly substitute:

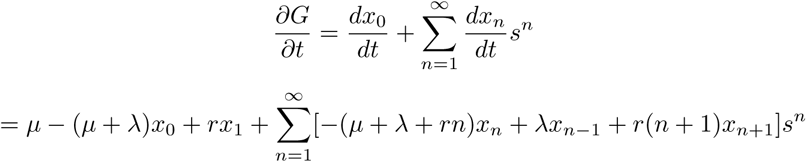

This simplifies to the following partial differential equation (PDE) for the probability generating function (PGF) G:

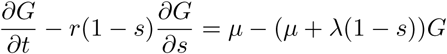

Because G is a probability generating function, a natural boundary condition is imposed wherein lim_s→1_*-G*(*t, s*) = 1 owing to the fact that the sum of the coefficients of the power series is equal to the sum of all the probabilities.

### 2.2 Statics

Note that if we let *µ* = 0 then we obtain the equation whose solution is the generating function of a Poisson distribution, aligning with known results [2]. Including demography or treatment forces the stationary distribution to belong to a different class. To find this class, we will start by solving for the stationary distribution whose generating function can be found by setting 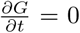, leading to the following ODE:

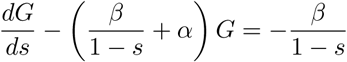

with the condition that lim_s→1_*-G* = 1 which is true for any probability generating function. In the static case, the stationary distribution is parameterized by two parameters: *α* = *λ/r* corresponding to the expected number of infections per recovery, and *β* = *µ/r* which is the expected number of demographic events or treatments per recovery. As the parameters are independent of s, this equation can be solved with the application of an integrating factor:

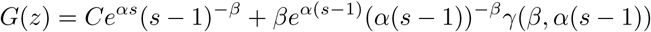

where *γ*(*a, x*) is the lower incomplete gamma function. Because any PGF has the property that the left limit as *s* approaches 1 should equal 1 and the term next to the integration constant has a singularity at 1, the constant must be zero. Therefore the PGF for the steady state distribution of MOI is given by

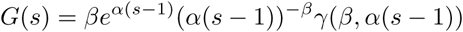

Replacing *γ* with its power series representation yields

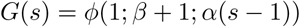

where *ϕ*(*a*; *b*; *x*) is Kummer’s confluent hypergeometric function, defined by

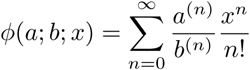

where *y*^(n)^ is the rising factorial of *y*. This is the PGF of the alternative hyper-Poisson distribution, *AHP* (*α, β* + 1) [7], whose probability mass function (PMF) is

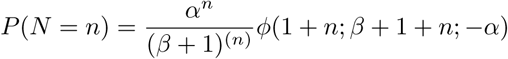

This distribution is a more general form of a Poisson distribution with *β* acting as a dispersal parameter. As *β >* 0 for any model that includes demography or treatment, the distribution is overdispersed compared to a Poisson distribution. A plot of the distribution for varying *β* is given in Figure 2. Note that despite qualitative similarities, it is distinct from the negative binomial distribution in both form and derivation - and in many cases due to its similarity to the Poisson distribution, as it is in the same family, it is sometimes a more appropriate alternative overdispersed distribution.

**Figure 2:**
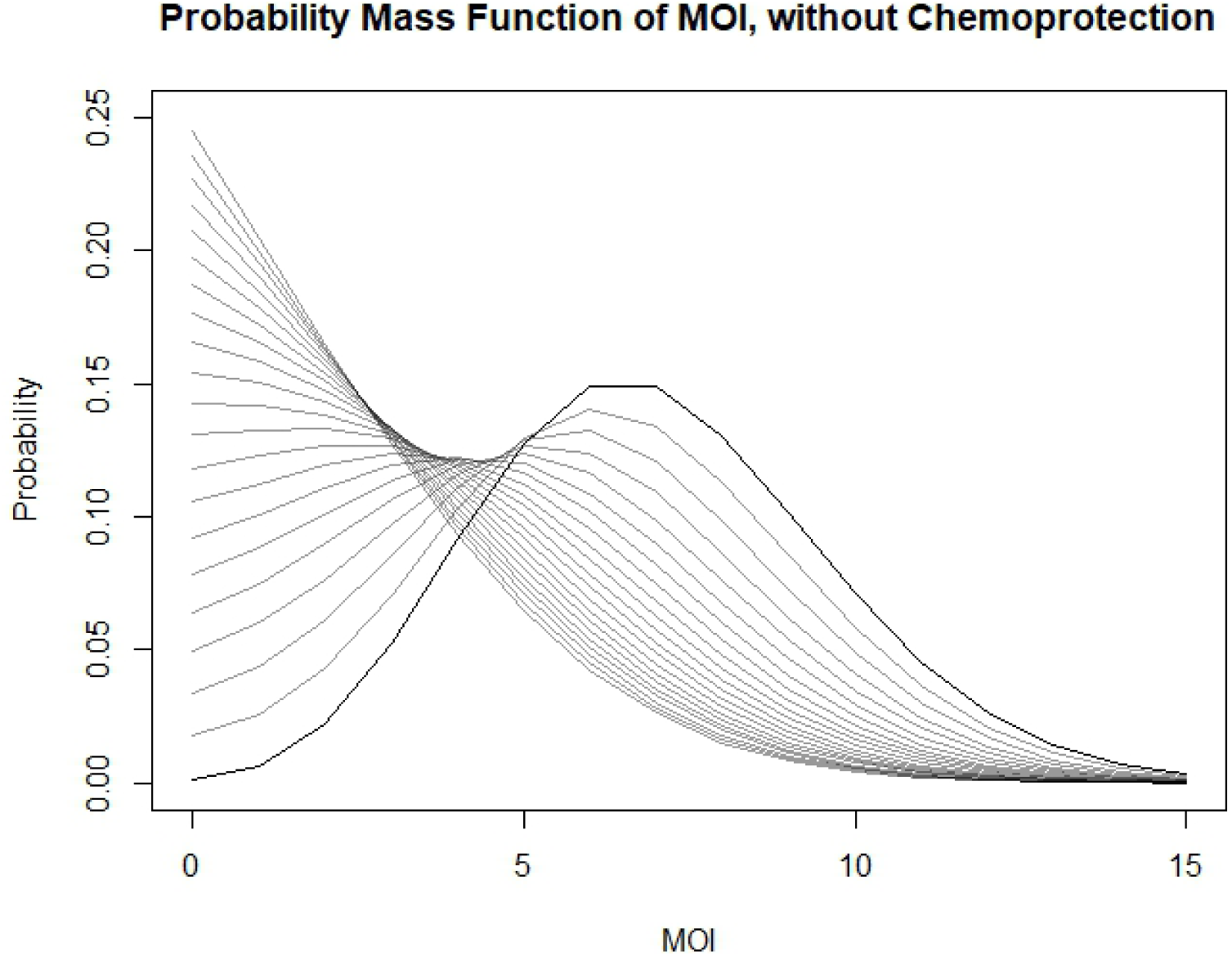
Plots of the PMF of the stationary distribution of MOI with varying values of treatment and demography, corresponding to the alternative hyper-Poisson (AHP) distribution. *α*, which is proportional to the force of infection, is held constant at 7. We vary values of *β*, defined here to be the ratio of the rate of treatment or death to the rate of recovery. *β* is varied from 0 to 2 in increments of .1. The bold line represents a Poisson distribution, *β* = 0. Increasing *β* suppresses MOI and increases the probability of having fewer infections, and it increasingly resembles a geometric distribution

Knowing the distribution of MOI is sufficient to get an estimate of the expected prevalence. Prevalence in this model is the complement of the proportion of individuals with zero infections, which is equal to the PGF evaluated at zero. Figure 3 shows plot of the prevalence as a function of the force of infection under varying treatment rates using this conversion. As expected, prevalence increases with FOI and decreases with treatment.

**Figure 3:**
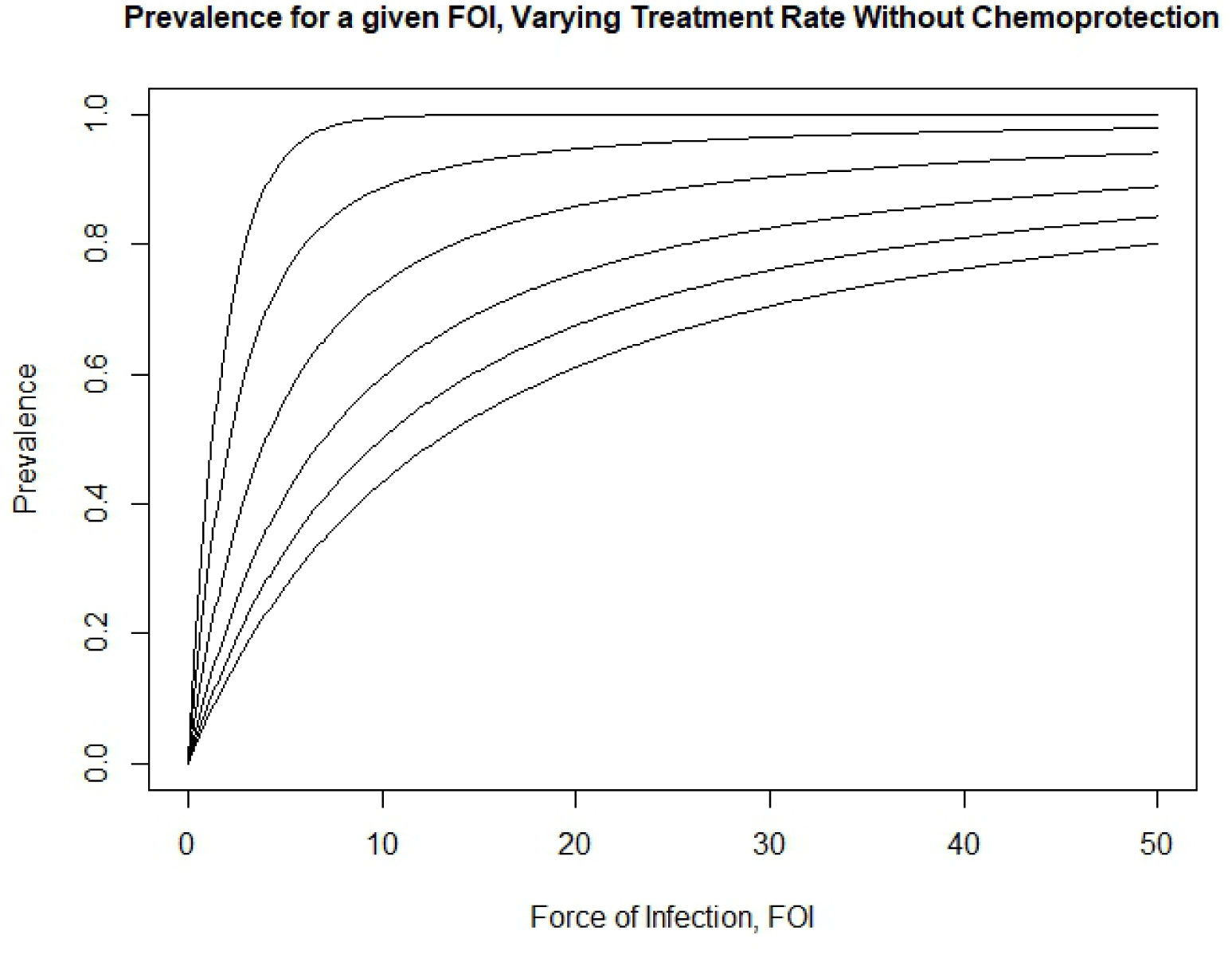
Plot of the prevalence as a function of annual FOI, excluding short term protection from treatment. The curves represent the expected prevalence for increasing FOI with set treatment rates. Prevalence monotonically decreases with increasing treatment rate. From top to bottom, the curves represent treatment rates of 0, 1, 3, 6, 9, and 12 per year.

**Figure 4:**
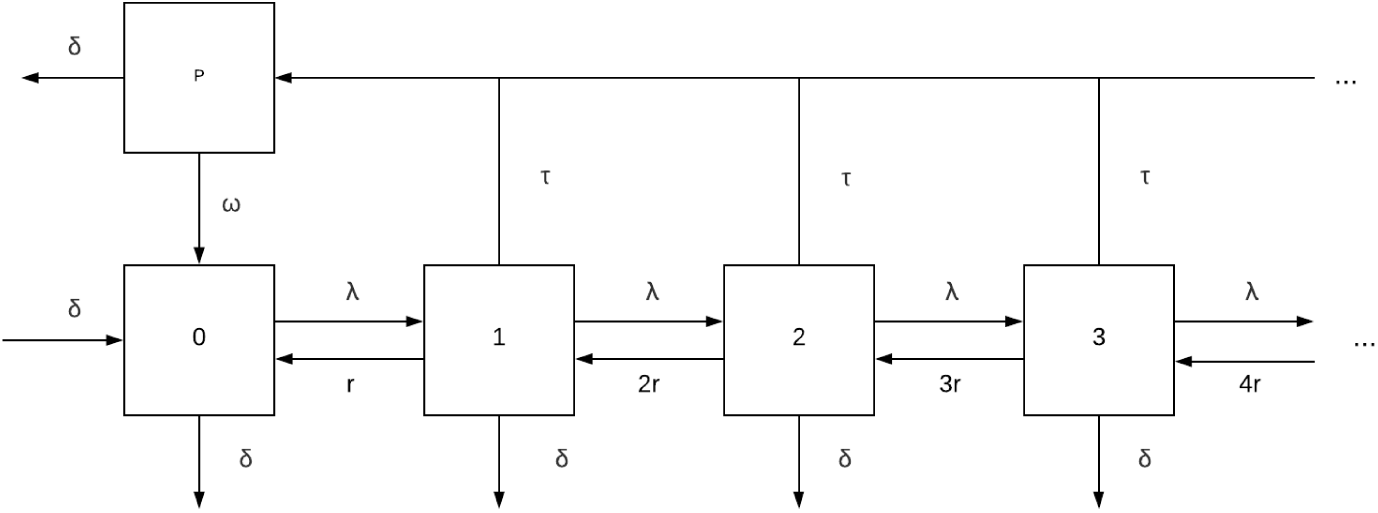
State space of human population with chemoprotection. Numbers indicate the number of infections in an individual and the letters denote transition rates between states, and the *p* compartment denotes the protected class

### 2.3 Dynamics

Now that we know the stationary distribution, we can turn our attention to the dynamics of MOI. In order to proceed, note that the transient distribution will always be AHP-distributed with *α* and *β* as being (possibly time-varying) parameters for the distribution. As these parameters depend on the three original parameters *λ*, *r*, and *µ*, they satisfy their own ODEs which will be derived here. This invariance of the class of AHP distributions under the dynamics can be seen by noting the set of PGFs corresponding to the class of AHP distributions constitute the only nontrivial solutions to the PDE, and once the system has entered this class it will not leave it without external perturbation.

To begin we go back to our PDE for the PGF and take a partial derivative with respect to *s*:

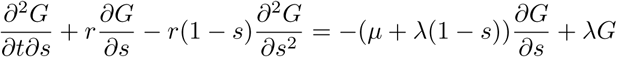

Plugging in *s* = 1 gives us the governing equation for the first factorial moment, also equal to the first raw moment:

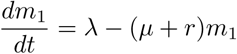

we then repeat the previous process with a second partial derivative to find an equation for the second factorial moment:

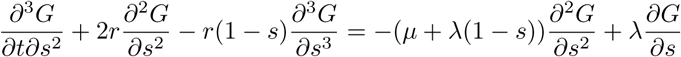

from which we can derive the equation for the second raw moment

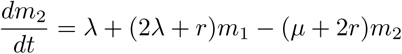

using the relationship between the second raw moment and the second factorial moment, that is

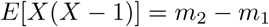

We then can match parameters to moments to determine the relationship between the transient moments and the transient parameter values that codetermine the exact distribution.

For this we note that for a random variable *X ∼ AHP* (*α, β* + 1),

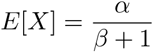

and the variance is

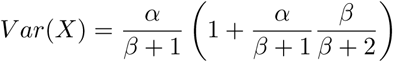

Matching moments allows us to get a functional relationship between the dynamics of the moments and the shape of the distribution. Doing so gives us

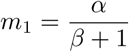

and

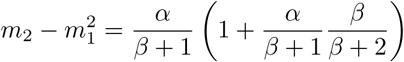

solving the system in terms of *α* and *β* gives us

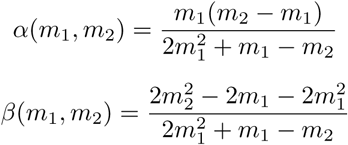

In particular, if we assume the system reached the limiting PMF at some point in the past without subsequent perturbation, then this combined with the PMF of the AHP distribution gives us a method of modeling the prevalence with a much more manageable system of two ODEs for the two moments necessary to parameterize the transient distribution:

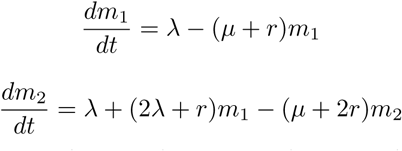

which we can convert back to prevalence *x* by noting that prevalence is the complement of the probability of having zero infections:

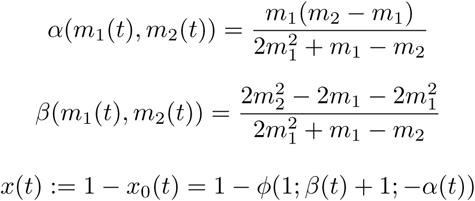

Note that we can also find *x*_n_(*t*), the proportion of individuals with *n* infections, by considering the PMF of the stationary distribution:

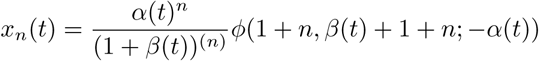

with the same definitions as before.

In addition, this means that if we are given an explicit time-dependent functional form for the FOI *λ*, we would be able to find the closed form solution for prevalence over time by solving the inhomogeneous system of two linear equations for the moments iteratively through the use of integrating factors. That is, in the language of queueing theory, we could find the transient solution for the PMF of states given the initial distribution is AHP-distributed. This assumption will be relaxed and a more general argument for the behavior of solutions will be made in the next section.

## 3 Stability of the AHP Distribution

Now that we know the stationary distribution, we can check its stability in the set of distributions. That is, if we don’t start off initially with an AHP distribution or it is perturbed through a non-uniformly acting intervention such as a targeted treatment or vaccine, will the distribution of MOI return to being AHP-distributed? If so, the dynamics above which assume AHP-distributed MOI will be good long-term approximations for any initial distribution. To investigate this, consider the PDE for the PGF:

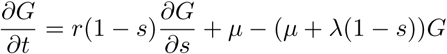

As this is a linear equation, we see we can arrange it to be in the following form:

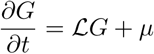

Where

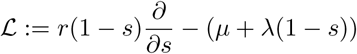

We will restrict the domain for *ℒ* to the following:

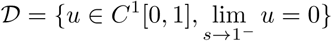

It’s clear that this forms a complete subspace of *C*^1^[0, 1] under the *L*^2^ norm.

Now we can define

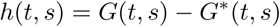

where *G* is the generating function for the transient distribution at time t and *G*^∗^ is the generating function for the stationary distribution found before. In particular this means 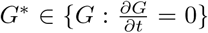 As the left hand limit of each of the functions on the right at 1 is 1 because they are PGFs, the left hand limit of *h* must be 0. Also, as long as the mean of *G* is finite, *h* will be once continuously differentiable. Therefore, *h ∈ 𝒟*

Now that we know h is in our domain, we can consider the action of 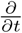:

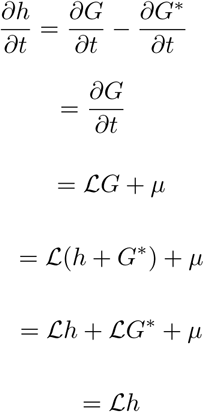

where the last equality holds because *G*^∗^ being stationary implies that ℒ *G*^∗^ = *-µ*. Now we can investigate the spectrum of ℒ *| 𝒟*, that is the operator restricted to the subspace that h is in. Setting up the eigenvalue problem, for any eigenfunction *u ∈ 𝒟* we get

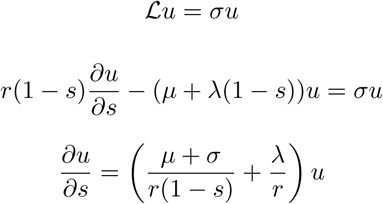

The parameters in the equation above for the stationary distribution are constant with respect tos. The solution to this equation is therefore

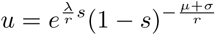

where the constant of integration is left out as it is irrelevant here to the span of the eigenfunctions.

The condition that the left hand limit at 1 always equals zero implies that the left hand limit at 1 of the time derivative is also zero. Because the action of the time derivative is equivalent to the action of the linear operator *ℒ* in the domain to which u belongs, this implies that the function can’t be singular at 1, and in particular

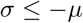

Therefore, the spectrum of *ℒ* is (*-∞,-µ*]. This combined with the expression for the time derivative of h yields

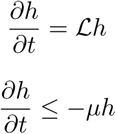

So

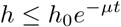

Plugging in the definition of h, we get

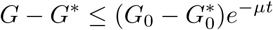

In addition, because PGFs in the unit interval are bounded by 1, *h*_0_ is bounded in absolute value by 1. Using this, we see

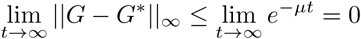

This implies uniform convergence of any initial PGF to that of an AHP distribution on the unit interval. Therefore any initial distribution with finite mean converges in distribution to an AHP distribution, so it is globally asymptotically stable in this domain as long as the parameters are non-negative and asymptotically constant with *r >* 0 for all time. For this system *µ* here acts as a sort of bifurcation parameter, where *µ* = 0 is the boundary between stability and instability. It is still stable at this boundary because any function which is not in the null space of the linear operator will have a strictly negative time derivative, which implies the asymptotic stability of the stationary distribution holds for the standard M/M/*∞* queue with its stationary Poisson distribution as well.

## 4 Model with Demography and Treatment, with Chemoprotection

Now we will consider the same system as before but we further assume that treatment confers some temporary protection from subsequent infection. That is, once someone is treated they enter a protected state as the treatment will linger in the body and protect against new infections for an average of 1*/ω* units of time before the individual returns to a susceptible state. The distribution of MOI conditioned on not being protected will remain AHP-distributed because adding this compartment does not change the queuing system - it adds a “waiting room” between bouts in the queue. Repeating the analysis done before by introducing a PGF results in another AHP-distribution, but with a zero inflation introduced by the proportion of individuals in the protected class. Furthermore, we can also repeat the stability analysis on the PDE/ODE system and see that it is also stable in the sense that any initial distribution with finite mean will asymptotically approach the zero-inflated AHP distribution. Treatment with waning protection therefore has three major effects on the distribution of MOI: decreasing the expected value, inducing overdispersion relative to a Poisson distribution, and zero-inflation. Each of these effects can individually correspond with an overall decrease in prevalence. See figures 5 and 6 for plots of the PMF of MOI and prevalence curves, respectively.

**Figure 5:**
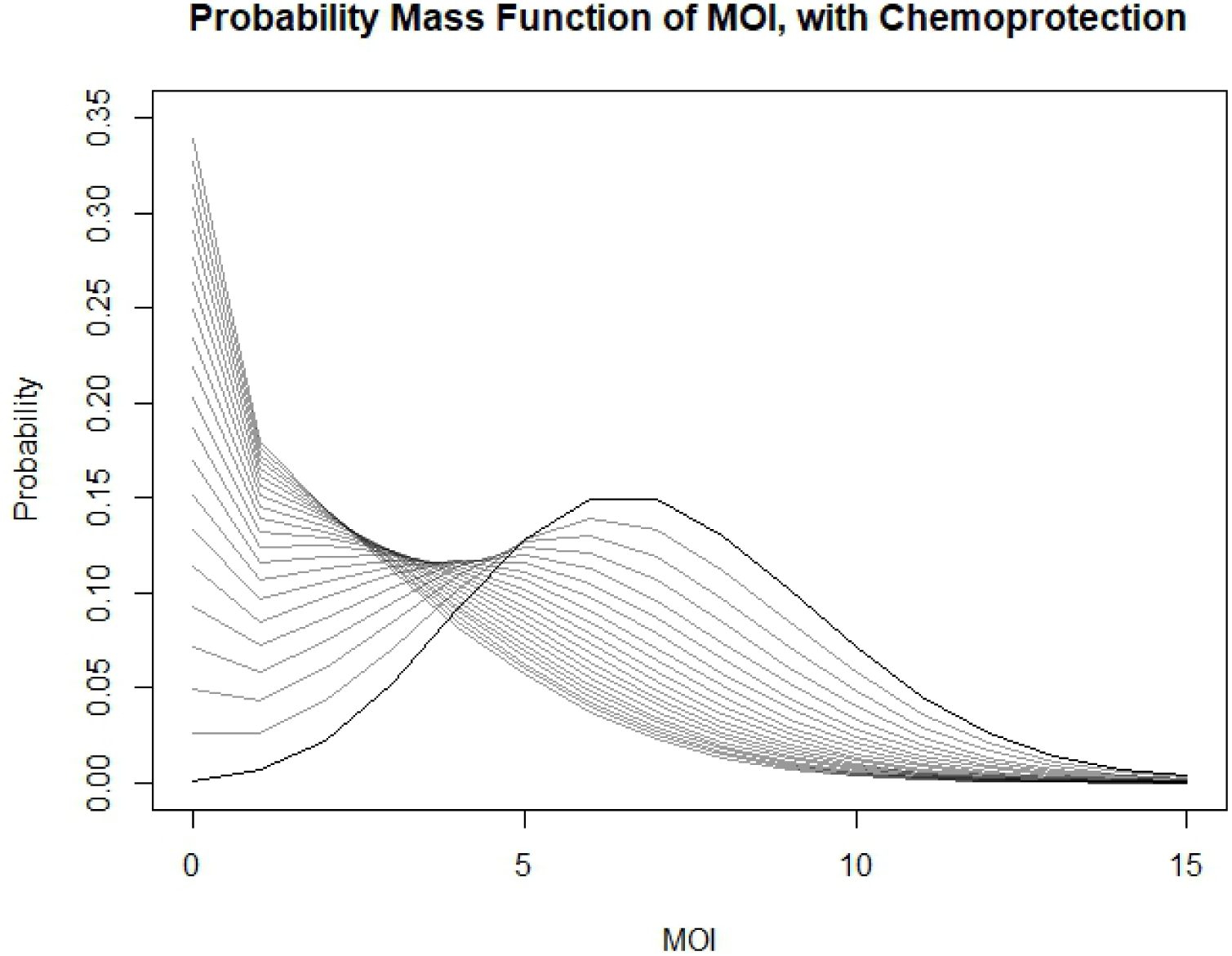
Plots of the PMF of the stationary distribution of MOI with varying values of treatment and demography, corresponding to the zero-inflated alternative hyper-Poisson (AHP) distribution. Note the distribution looks the same as in figure 2, but with increased probability mass at zero due to the fraction of the population in a protected state. Mean protection time (1*/ω*) is set to 30 days.

**Figure 6:**
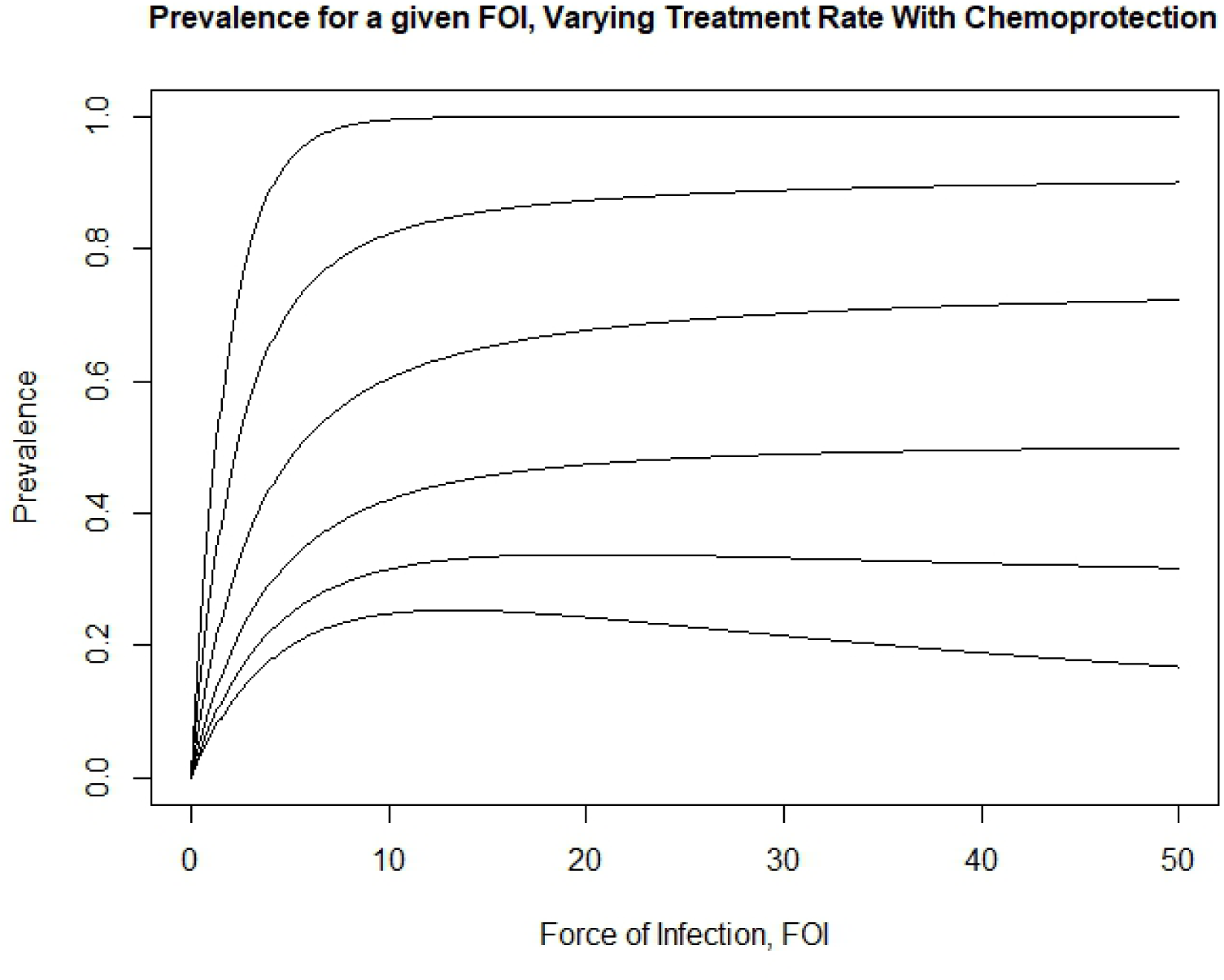
Plot of the prevalence as a function of annual FOI, including short term protection from treatment. Average duration of protection is set to 30 days. The curves represent the expected prevalence for increasing FOI with set treatment rates. Prevalence monotonically decreases with increasing treatment, but at high treatment rates prevalence will drop with increasing FOI due to the increase to the protected class. From top to bottom, the curves represent treatment rates of 0, 1, 3, 6, 9, and 12 per year.

With the same notation as before and letting *p* represent the proportion of individuals in the protected class, the full model with the protected state can be written as

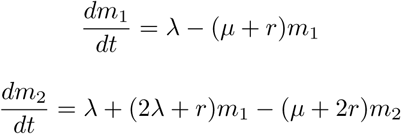

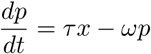

Where

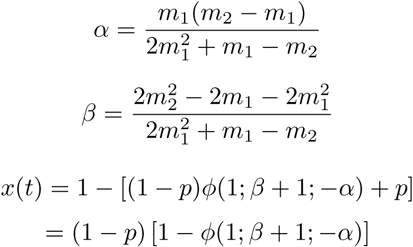

Note that *x* has a nice physical interpretation in this form - it is the probability that an individual has at least 1 infection given that they are not protected, times the probability they are not protected.

## 5 Pseudoequilibrium Model

Now that we have transient behavior established, we can go one step further and include a pseudoequilibrium analysis. That is, rather than having to deal with variables which parameterize the distribution we can assume the distribution approaches equilibrium on timescales that are much smaller than the rate at which the force of infection changes. This was implicitly made in the model developed for the Garki project [2], wherein the rate of recovery was given as the reciprocal of the expected duration of infection. If we denote the time spent infected as *X*, then the expression for the expectation is as follows:

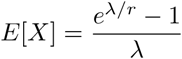

It’s useful to compare this to the case without superinfection. Because each infection has an exponentially-distributed time to clearance, without superinfection the expected time spent infected is 1*/r*. In the case with superinfection, we can see first that it is monotonically increasing with respect to the force of infection. In addition, for small FOI the Taylor expansion of the exponential shows that the expected time is approximately 1*/r*:

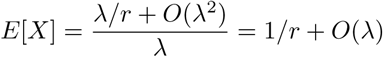

Now we can consider the same simplification with treatment and demography. In order to consider this case, we need to know how this expectation changes when introducing exponentially-distributed treatment and demography. Although the distribution of the time to recovery starting at a single infection is not known in closed form, its Laplace transform is - and we can exploit that to determine the impact of these perturbations [5]. Denoting the time until treatment (or death) with *T*, then the total duration of infection *D* is *D* = min {*X, T}* That is, the infection will end if an individual either recovers naturally or gets treated or dies, whichever comes first. Then we can write down the following:

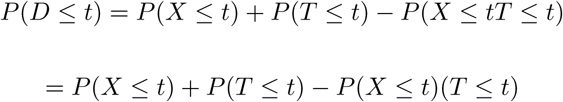

where this step uses the independence of the natural recovery and treatment/demography processes. As these are exactly the CDFs of the respective variables, we get

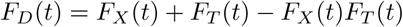

Because treatment and death are exponentially distributed, the minimum of the two is also exponentially distributed with a rate equal to the sum of their rates. We will again denote this new rate as *µ*. Knowing this allows us to plug in the CDF of this exponential distribution for *F_T_* :

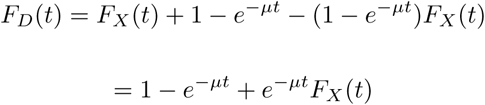

Using the fact that the integral of the complement of the CDF is equal to the expectation for any random variable, we get

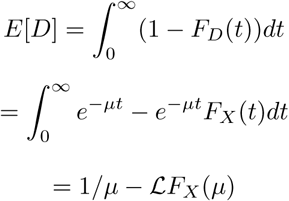

where here *ℒ* denotes the Laplace transform. Finally, using the relationship between the Laplace transform and the integral of a function gives us a formula for the expected duration of infection as a function of the Laplace transform of X:

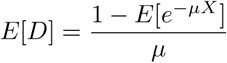

In proposition 4.1 from the paper of Guillemin et al, the Laplace transform of the duration spent infected (there called the “congestion duration”) starting with a single infection is given by

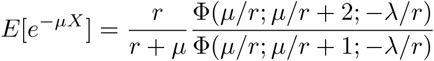

which, plugging in, gives us the expected duration of an infection with superinfection and treatment

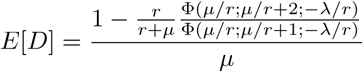

Although this rate looks complicated it is consistent; the limit as *µ* goes to zero is exactly the rate found previously and implemented in the Garki model [2].

Before plugging this in to a model of infection and recovery, we must also determine the proportion of people who were treated rather than died or recovered, as they will enter the protected class. This can by found again through a probabilistic argument:

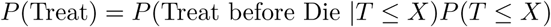

That is, the probability of being treated to end the infection is the product of the probabilities of ending the treatment through either treating or dying given that an individual won’t recover naturally, times the probability that the individual won’t recover naturally. The first probability is trivial; it’s the probability that one exponential random variable is less than another, and in this case is *τ/*(*τ* + *δ*) = *τ/µ*. The second probability is more interesting, since again with superinfection X is not exponentially-distributed. Again we can write this probability out:

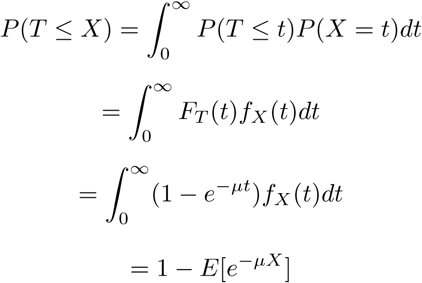

which shows that the Laplace transform of a random variable X can also be interpreted as the probability that a random variable defined over the nonnegative reals is less than or equal to an exponential random variable with rate *µ*. Note that this quantity as a probability is bounded between 0 and 1, and as *µ* increases to infinity this probability decreases to 0. Multiplying those probabilities, we see

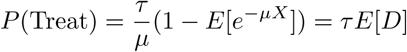

Therefore, we can complete the simplified model by replacing the simple rate of recovery r from infected to susceptible with the reciprocal of this duration of infection as follows:

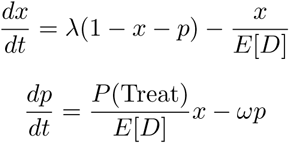

which we found can be written as

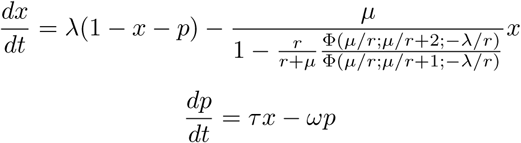

This reduces the model down to just the two equations that will adequately describe what proportion of individuals infected in a single variable, *x*, given that the distribution of MOI which determines the rate of recovery is transiently at (or very close to) its steady state.

## 6 Connecting to Ross-Macdonald Dynamics

Now that we have fully established the system of infections and recoveries on the human side, we can couple the human system with a vector system to observe the transmission dynamics. This is the hybridization step, wherein we reinterpret the probability distribution of MOI as a frequency distribution for a large population. This coupling with simple vector dynamics gives us a Ross-Macdonald style model. Ross-Macdonald models are SIS compartment models with vector-based transmission used to model malaria transmission. There is no single canonical form for a Ross-Macdonald model (see for example the review by Smith et al. for an overview of the history and main developments in Ross-Macdonald models [12]), but as an example we chose the one implemented by Smith and McKenzie because there is only one additional equation for mosquitoes without any delay dynamics, but the extrinsic incubation period is still taken into account [13].

In this model the force of infection is proportional to the number of infectious mosquitoes. That is,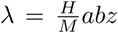 where 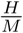 is the ratio of humans to mosquitoes, *a* is the rate at which mosquitoes bite humans, *b* is the proportion of infectious bites which result in human infections, and *z* is the infectious fraction of mosquitoes. Replacing this expression for the force of infection in the original system for the PGF of the MOI G, we get the following PDE/ODE system:

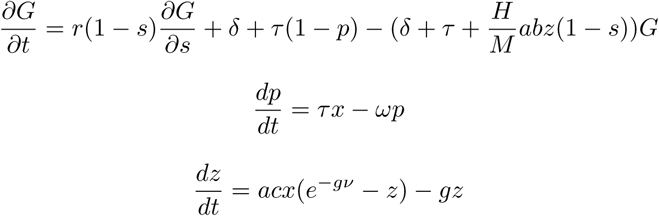

With

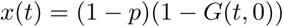

where *x* is the proportion of humans infected, *c* is the probability an infectious human will infect a susceptible mosquito during a blood meal, *g* is the death rate of mosquitoes, and *ν* is the extrinsic incubation period. As done before, we include the following boundary condition for the PGF G:lim_s→1_*-G*(*t, s*) = 1. This system will hold for any biologically reasonable initial distribution of MOI and fraction of infectious mosquitoes.

However if our initial MOI is AHP-distributed, then the dynamics can be entirely captured by a system of four ODEs:

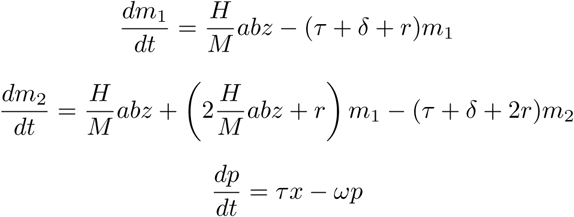

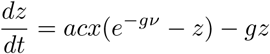

Where

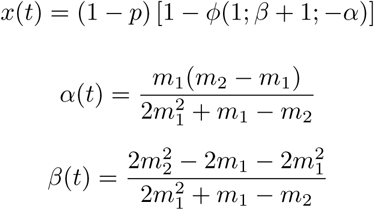

*c* is the probability an infectious human will infect a susceptible mosquito during a blood meal, *g* is the death rate of mosquitoes, and *ν* is the extrinsic incubation period. These ODEs are derived in the same manner as before, this time tracking the extra compartment.

This ODE system is a bit different from most implementations of the Ross-Macdonald model, but similar to the one introduced by Nåsell [9]. The first two equations determine the moments of the distribution of MOI while the third determines the degree of zero-inflation. These are converted to current prevalence by taking the complement of the probability an individual has zero infections. This in turn affects the rate of recruitment of mosquitoes to the infectious class in the fourth equation, which is assumed proportional to the force of infection acting on the distribution of susceptible people.

It’s important to recognize that this will hold if the initial distribution is in the null space of the linear operator described in the previous section, which corresponds to the class of AHP distributions. However, because we proved any distribution asymptotically approaches the class of AHP distributions, knowing malaria has been endemic in a region for a long period of time without strong perturbation to the structure of the system is enough to make this a reasonable assumption for modeling local transmission.

Finally, in the specific case where there is no lasting protection from treatment we can also couple the pseudoequilibrium simplification to transmission dynamics in an analogous way to achieve a set of three simpler, albeit still nonlinear, ODEs:

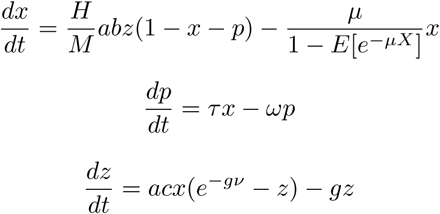

Where

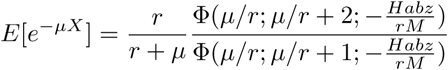

The interpretation of this formulation is much more straightforward than the previous two versions - we no longer need information about the entire distribution. In particular, this form makes it clear that the distribution of MOI only affects the rate of recovery and it is therefore much more directly comparable to other Ross-Macdonald models. However, it is important to remember this model only holds when the transient distribution is exactly equal to its stationary distribution at every point in time.

## 7 Discussion

In this paper we introduced a model of malaria superinfection which properly accounts for treatment and demography, two epidemiologically-relevant features of malaria which influence the distribution of MOI and have nonlinear effects on the mean recovery rate. As the proportion of the human population with a single infection represents the only group which can recover naturally at a given time, these perturbations to the recovery rate can propagate through to differentially affect the modeled effect size of drugs on transmission. In particular, this effect will be highly context-dependent - a drug will have a stronger measured effect on prevalence in areas with a high force of infection than predicted in a model without superinfection.

In addition, temporary protection conferred by drugs can have a major impact on the prevalence,as shown in comparing figures 3 and 6. This “waiting room” for the infection queue has the effect of pulling down the equilibrium prevalence, and nonlinearly impacting the influence of FOI on prevalence. This has interesting and somewhat troubling implications for malaria control in areas with high treatment rates - reducing the effective FOI through any intervention appears to has the opposite intended effect on overall prevalence.

While this model represents a substantial advance and generalization of past results, it is important to keep in mind the underlying assumptions when interpreting the model predictions in real transmission systems. Because infection events are modeled as a simple Poisson process, an infectious bite can only result in a single infection. This allows the original M/M/*∞* model to be “tridiagonal” in the sense that an individual can only increase or decrease the number of infections by one at a time. However in areas of high transmission intensity, a bite from a superinfected mosquito may establish multiple infections from the same blood feeding event [1]. Parasite diversity can also contribute to transmission; for example, deletion of HRP-2, the target of common rapid diagnostic tests, can decrease the rate of detection and subsequent treatment of infections [6]. There is also evidence to suggest a host who becomes anemic due to periods of high parasitemia may be less prone to infection due to the efflux of iron from the liver [11]. All of these break Macdonald’s original assumption of independence of the infections, and generalizations which account for these could have important implications for malaria control and elimination strategies. Future explorations of these factors is needed.

Despite these caveats, this model serves two main purposes. First, it provides a consistent framework with respect to the underlying assumptions of the stochastic process for investigating the impact of drugs and demographic turnover on the prevalence of any pathogen which can superinfect its host. Second, it provides an analytic demonstration of how the distribution of MOI under the effects of superinfection is inextricably connected to the structure of the rest of the model, as well as some tools for investigating the impact of this kind of heterogeneity. This connection and the dynamics that it generates could change the policy recommendation for intervention packages in many regions, impacting the lives of all people living in endemic areas. Because of that, we must be cognizant of the impact of modifications of this type of hybrid model on the structure of the distribution of MOI as we include more detail and realism in our attempts to better understand and prevent the spread of infectious disease.

## Acknowledgements

This work was made possible through grant OPP1110495 from the Bill and Melinda Gates Foundation.

I would like to acknowledge and thank David L. Smith, Daniel Citron, Sean Wu, and the rest of the MASH malaria team for their comments and insights in preparing this manuscript.

